# Contrast and luminance gain control in the macaque’s lateral geniculate nucleus

**DOI:** 10.1101/2022.08.29.505733

**Authors:** R.T. Raghavan, Jenna G. Kelly, J. Michael Hasse, Paul G. Levy, Michael J. Hawken, J. Anthony Movshon

**Affiliations:** Center for Neural Science, New York University, New York, New York, 10003

**Keywords:** LGN, luminance, contrast, temporal frequency, gain control

## Abstract

In natural scenes, there is substantial variation in the mean and fluctuation of light levels (luminance and contrast). Retinal ganglion cells maintain their sensitivity despite this variation and their limited signaling bandwidth using two adaptive mechanisms, which control luminance and contrast gain. However, the signature of each mechanism and their interactions further downstream of the retina are unknown. We recorded neurons in the magnocellular and parvocellular layers of the lateral geniculate nucleus (LGN) in anesthetized adult male macaques and characterized how they adapt to changes in contrast and luminance. As contrast increases, neurons in the magnocellular layers maintain sensitivity to high temporal frequency stimuli but attenuate sensitivity to low temporal-frequency stimuli. Neurons in the parvocellular layers do not adapt to changes in contrast. As luminance increases, magnocellular and parvocellular cells increase their sensitivity to high temporal frequency stimuli. Adaptation to luminance is independent of adaptation to contrast, as previously reported for LGN neurons in the cat. Our results are similar to those previously reported for macaque retinal ganglion cells, suggesting that adaptation to luminance and contrast result from two independent mechanisms that are retinal in origin.

## Introduction

There is substantial variation in both mean light levels (luminance) and their local fluctuation (contrast) in natural scenes (Frazor and Geisler, 2006; Webster and Mollon, 1997). Two mechanisms allow the early visual system to adjust to these changes, gain controls for luminance and contrast (Shapley and Enroth-Cugell, 1984). Each mechanism alters the firing of visual neurons to preserve information about the pattern of light on the retina, given wide variations in contrast and luminance and limited neuronal firing rates. How each adaptation mechanism influences a cell’s sensitivity can differ based on cell class. For the two major ganglion cell classes of the cat retina, for example, contrast gain control exerts a more considerable impact on the responses of Y cells than X cells (Shapley and Victor, 1978). In addition, Y cells become light-adapted and transient at lower luminance levels than X cells (Jakiela et al., 1976). Similar distinctions hold for the ganglion cells that project to the magnocellular (M) and parvocellular layers of the primate lateral geniculate nucleus (LGN) (which we will term M and P ganglion cells), as well as activity for the cells within these layers (which we will term M and P cells). Contrast gain control has more effect on M ganglion cell responses (Benardete et al., 1992; Benardete and Kaplan, 1999). Moreover, as in cat Y cells, M ganglion cells become light-adapted and transient at lower light levels than P ganglion cells (Purpura et al., 1990).

M and P pathway cells also differ in ways that X and Y pathway cells do not. Response gain, the initial slope of firing rate as a function of contrast, is higher in the M pathways than the P pathways for ganglion cells in the retina and recipient M- and P-layers in the LGN (Derrington and Lennie, 1984; Kaplan and Shapley, 1986; Movshon et al., 2005). Moreover, while M ganglion cells maintain a significant response gain at scotopic light levels, P ganglion cells have negligible response gain at these light levels (Purpura et al., 1988). Neither of these properties distinguishes X from Y cells (Shapley, 1992; Shapley and Enroth-Cugell, 1984). However, within the X and Y cell classes, there is a clear relationship between the size of the center receptive field mechanism and the strength of luminance and contrast adaptation. As the center size increases, the effective luminous flux (area × intensity) over the center increases, and cells become light adapted at lower light levels and have higher contrast gain (Enroth-Cugell and Shapley, 1973; Shapley and Enroth-Cugell, 1984). Differences in receptive field size might account for the differences in contrast gain between M and P ganglion cells (Purpura et al., 1988). M ganglion cells have larger dendritic fields than P ganglion cells (Dacey and Petersen, 1992), and measurements of the receptive field (RF) center in both the retina and LGN have shown that M ganglion cells and M-layer LGN cells have a larger RF center than retinal and LGN P-layer cells (De Monasterio and Gouras, 1975; Derrington and Lennie, 1984). Changes in filtering properties parallel the changes in response dynamics caused by luminance and contrast gain control. As contrast levels decrease, cat X and Y cells and monkey M ganglion cells become lowpass temporal filters (Benardete and Kaplan, 1999; Shapley and Victor, 1978).

On the other hand, P ganglion cells exhibit little change in their temporal frequency selectivity with changes in contrast (Benardete et al., 1992), suggesting that contrast gain control does not significantly impact this pathway. As light levels increase, both M and P ganglion cells exhibit a similar elevation in response gain at high temporal frequencies (Purpura et al., 1990). There appears to be little interaction between the luminance adaptation and contrast gain control for cat X and Y LGN cells at photopic light levels, as changes in luminance and contrast exert separable effects on these cells’ temporal frequency tuning (Mante et al., 2005). It remains unclear whether the same holds in the primate M and P system.

Here we seek to address two gaps in our knowledge about how neurons in the M- and P-layers of the monkey LGN adapt to luminance and contrast. First, we wanted to know whether LGN neurons adapt their response in the same ways reported for retinal ganglion cells that project to the M- and P-layers (Kaplan and Shapley, 1984; Lee et al., 1990). While some studies suggest that short-term adaptation of LGN responses matches the signatures of retinogeniculate cells (Spekreijse et al., 1971; Virsu and Lee, 1983), there is also evidence that the thalamus strengthens the contrast gain control carried by retinogeniculate inputs (Alitto et al., 2019; Kaplan et al., 1987). Second, we wanted to test whether the independent luminance and contrast gain controls observed in cat X and Y pathways also exist in the primate. Understanding the gain control signatures of the M and P pathways at this level of detail is critical to developing models of visual representations in the retina and thalamus. Moreover, given that the M and P pathways remain partially segregated from the input layers of the primary visual cortex to the early extrastriate cortices (Callaway, 2005; Sincich and Horton, 2005), any complete account of cortical visual representations may need to take into account subcortical gain mechanisms within the M and P pathways. Our findings demonstrate that LGN neurons seem to faithfully represent known changes in retinal input under variations in luminance and contrast, that changes in contrast and luminance exert separable influences upon the temporal sensitivity of macaque LGN neurons, and that the influence of each adaptation mechanism varies as a function of both cell class and eccentricity.

## Methods

We recorded three visually normal male macaques, a 9-year-old *Macaca nemestrina*, an 11-year-old *Macaca mulatta*, and a 3-year-old *Macaca fascicularis*. We used an anesthetized paralyzed preparation described in detail previously (Movshon et al., 2005). Briefly, we induced anesthesia with an intramuscular injection of ketamine HCI (10 mg/kg) and maintained anesthesia during catheterization of saphenous veins and endotracheal intubation with isoflurane. After placement of the animal into a stereotaxic frame, we maintained anesthesia with an infusion of between 6 and 30 μg/kg/hr of sufentanil citrate and neuromuscular blockade with an infusion of 0.1 mg/kg/hr vecuronium bromide to limit eye movements. We opened a craniotomy and durotomy to allow a 23G guide tube containing a glass-coated tungsten microelectrode (Merrill and Ainsworth, 1972) to be lowered to a position 5 mm above the LGN; we then advanced the electrode out of the guide needle into the nucleus. Entry to the LGN was indicated by the onset of brisk time-locked visual activity in response to alternating red-green flicker. In most instances, we assigned recordings to layers of LGN based on the sequence of eye dominance across depths relative to the entry into the LGN, aided by functional criteria (described below). We made electrolytic lesions at points of interest along the microelectrode track for most penetrations. At the conclusion of the experiment, we euthanized the animal with an overdose of barbiturate (pentobarbital, 65 mg/kg) and perfused it with 4% paraformaldehyde in 0.1 M PBS. We took the brain and cut parasagittal sections at 50 μm on a freezing microtome. We reconstructed the electrode tracks based on tissue marks and electrolytic lesions visualized in Nissl stained sections to determine a laminar assignment for each recording site.

### Visual stimulation

We administered atropine sulfate (1%) drops to each eye to dilate the pupils (typical diameter 6 mm) and paralyze accommodation and placed gas-permeable +2D contact lenses in each eye to protect the corneas. The contact lenses were removed for cleaning each day. We estimated refractive error by direct ophthalmoscopy and refined the estimate by optimizing the response of visual units. We chose supplementary spectacle lenses to make the retinas conjugate with a screen 114 cm distant.

We generated and controlled stimuli with an Apple Mac Pro computer. We presented stimuli on a CRT monitor (HP1190) running at a resolution of 1280 × 960 pixels (64 pixels per degree) and 120 Hz. Most stimuli were drifting sinusoidal gratings whose mean luminance took values between 3.5 and 41 cd/m^2^, yielding an estimated retinal illuminance of 100-1150 Td.

We located the center of each LGN neuron’s receptive field with hand-controlled targets. We then measured each cell’s contrast response, spatial and temporal frequency tuning, chromatic selectivity, and size tuning in separate experiments in which only one of these parameters varied while the others were fixed. In the main experiment, we recorded the response of each neuron to large-field (20×15 degrees of visual angle) sinusoidal drifting gratings of a spatial frequency near each cell’s optimum (typically around 1 c/deg). We modulated the temporal frequency of these gratings over time – it either increased smoothly exponentially over 14 seconds from 0.5 to 32 Hz or increased in equal ratio steps every 1.6 seconds from 0.5-40 Hz. We refer to the first stimulus as a *chirp sweep* and the latter stimulus as a *stepped sweep*. We presented each sweep at 3 luminance levels and 4 or 5 contrast levels.

Each block began with a blank screen of a given luminance displayed for between 1–1.6 seconds to allow cells to adapt. Five blocks of sweeps were run consecutively at the same mean luminance level, with contrast increasing by 0.5-1.0 octaves (a factor of √2 - 2) over the stimulus presentation interval of 14-24 seconds. Following the completion of 5 blocks, the mean luminance increased or decreased in the next block, and we made another set of contrast response measurements. We presented 4-5 contrast levels for each of the 3 mean luminances tested, with 5 repeats per block. We used contrasts between 0.0125 and 0.8 and luminances between 3.5 and 41 cd/m^2^ (estimated retinal illuminance 100–1150 Td) based on the range of stimulus luminance offered by our CRT display monitor and the precision offered by the bit depth of the display controller. At the lowest luminance, contrasts were between 0.05 and 0.8; at the highest luminance, contrasts were between 0.0125 and 0.4.

### Fitting LN model

We computed averaged response histograms, and low pass filtered them with a 2nd order Butterworth filter with a 40 or 60 Hz cutoff depending on the stimulus. For chirp stimuli, the high pass cutoff was 40 Hz, and for stepped stimuli, 60 Hz, to ensure that neural modulations at the highest temporal frequencies tested (32 and 40 Hz, respectively) were not attenuated. We used the full time course of the response for each combination of contrast and luminance to fit a linear-nonlinear (LN) model and used these models to estimate the response of each cell to temporal frequency as a function of contrast and luminance.

### Model components

The linear component of the model was a cascade of high and low pass filters introduced by Shapley and Victor (1981). In the frequency domain, this model takes the form

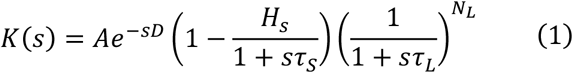

where *A* is the amplitude, *D* is an initial delay, *τ*_*s*_ and *τ*_*L*_ are time constants of the low and high pass filters, *H*_*S*_ is the high pass strength, and *N*_*L*_ is the number of stages in the lowpass filter, and *s* = *i ω*, where *ω* is the temporal frequency. The first term of this equation is the effect of a temporal delay in the frequency domain, while the second and third terms correspond to a high and low pass filter, respectively.

We multiplied the linear model by the stimulus *S* in the frequency domain, *S*(*ω*) ⋅ *K*(*ω*), and then inverse Fourier transformed the result to yield the filter output *k*(*t*) for a given set of model parameters. We refer to this filter output as the generator signal following Chichilnisky and Kalmar (2002), and it can be thought of as an approximation of the stimulus-driven membrane voltage. The generator signal *k*(*t*) passes through a nonlinearity to generate spike rate over time *r*(*t*). In this analysis, we used a rectifying nonlinearity defined as follows:

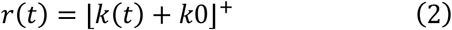

where ⌊ ⌋^+^ indicates rectification so that

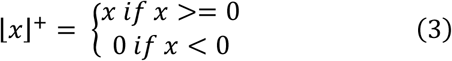

and *k*_0_ is a scalar representing the difference between the resting potential *V*_rest_ and the threshold membrane potential *V*_thr_ (Mechler and Ringach, 2002). Positive values of *k*_0_ allow the model to have spontaneous firing rates in the absence of visual stimulation. In control analyses, we introduced an exponent on this nonlinearity, but in almost every case tested, the best fitting exponent was close to 1. Since including an exponent as a parameter did not improve the fit substantially, we used the simpler form. The linear and nonlinear components of the LN model were fit separately.

### Fitting procedure for the nonlinearity

We assumed a fixed nonlinearity at each luminance condition estimated as follows. We measured the baseline firing rate at zero contrast for each luminance level. Preceding each run of the sweep stimuli described above, we presented a zero contrast blank screen at one of 3 luminance levels for 1-1.6 seconds. We gathered 25-40 seconds of zero contrast data per luminance level across 5 repeats and 5 contrast conditions. We formed a distribution of the binned firing rates at each luminance level. We assumed that the distribution of firing rates observed at zero contrast is the result of rectifying *k*_0_ when *k*(*t*) = 0. Prior work has shown that a rectified Gaussian can approximate this distribution of firing rates. The mean of this distribution represents *V*_rest_ − *V*_thr_, or *k*_0_. (Carandini, 2004; Mante et al., 2005). We estimated the mean and standard deviation of a rectified Gaussian distribution that minimized the negative log-likelihood of observing the distribution firing rates at zero contrast. This calculation was performed separately at each luminance level (three nonlinearities total), and *k*_0_ was set equal to the estimated mean.

### Fitting procedure for the linear filter

Across stimulus conditions, the fitting procedure allowed all linear filter parameters to vary. We used the linear filter parameters for M and P cells published in Benardete and Kaplan (1999, 1997) to set bounds on these parameters. We fit the LN model to both the time-varying mean firing rate and the mean of the measured first harmonic (F1) responses in each condition for both stepped and chirp sweep stimuli. The F1 is the amplitude and phase of the fluctuations in firing rate at the temporal frequency of the stimulus. The first harmonic response of a rectified linear filter (LN) model is derived as follows according to Mechler and Ringach (2002). Given the difference between the resting membrane potential and the threshold membrane potential *V*_rest_ − *V*_thr_ = *k*_0_ and the amplitude response of a linear filter |*K*(*ω*)|, the first harmonic response is calculated from the ratio

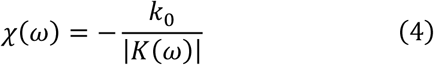

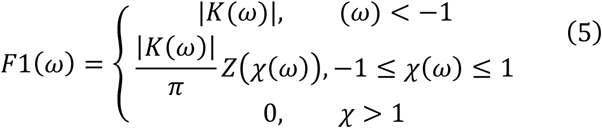

where

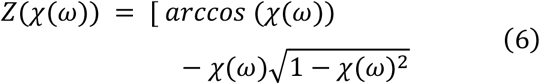

After calculating the F1 at each temporal frequency, we optimized filter parameters to minimize the sum of squared errors between the F1 response (amplitude and phase) of the model calculated using equation **(1)** and the measured F1 of the original data. We weighted the sum of squared errors using the square root of the amplitude response at each temporal frequency. Phase differences have a smaller magnitude than amplitude differences, so we reduced amplitude errors to bring phase and amplitude errors into roughly the same range and let them contribute more equitably to the model fit.

For chirp sweeps, the F1 could not be calculated below 4 Hz. Therefore, we optimized filter parameters to minimize the squared error between the time-varying response and the model prediction in time, weighted by the square root of the firing rate at each point in time. The fitting procedure was additionally constrained to match the F1 response that could be calculated at or above 4 Hz. We found this useful to prevent the model from overestimating the response of cells to high temporal frequencies where noise dominates the response for many cells.

### Descriptive model fitting

Below we describe in detail a set of models whose parameters were fit to a cell’s F1 response to spatial frequency, temporal frequency, and contrast. Model fitting in each instance was performed using standard bounded nonlinear optimization (interior-point algorithm). The cost function was defined as a weighted sum of square differences between the model predictions and data, where the weights – on the assumption of Poisson variability – were usually the inverse of the square root of the response at each condition.

### Separability analysis

We followed the methods of Mante et al. (2005) and used singular value decomposition to examine whether we could decompose contrast- and luminance-evoked changes in temporal frequency tuning into separable factors. We started with the complex output (both amplitude and phase) of the set of linear filters fit to data gathered at various luminance *L* and contrast *C* conditions. We can write the response at each frequency as *F*_*L,C*_ (*ω*), which is a complex matrix that is *LxC* conditions in size. We used singular value decomposition to partition this matrix into the form *USV*′, where the columns of U and V define vectors, each of which is a contrast and luminance-defined cross-section, the product of which yields a separable transfer function that approximates *F*_*L,C*_ at a given omega value. The degree to which a particular pair of columns approximates the data in *F* are given by the magnitude of the diagonal values of *S*. The magnitude of the first diagonal value in *S* provides an estimate of how well the function *F*_*L,C*_ can be approximated by a single pair of functions. A separability index can be defined (Depireux et al., 2001) as follows.

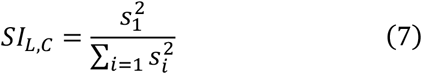

This index is between 0 and 1, with values closer to one being better approximated by a pair of separable functions. Analysis was usually performed on temporal frequency tuning curves derived from model fits. We also used the SVD directly on the measured F1 response as a control.

Two factors limit the efficacy of SVD in our current work. Since stimulus discretization and monitor gamut limit the contrast range that can be presented, stimulus contrasts lower than 0.05 or higher than 0.4 could not be rendered at every luminance level presented. SVD cannot operate on a matrix with incomplete or missing entries. For experiments on P cells, this typically means we operate the SVD on contrast conditions between 0.05–0.4, while for M cells, we examined contrast conditions between 0.05–0.2, for which we could make measurements at every combination of contrast, luminance, and temporal frequency. A second limitation is that occasionally a contrast level presented is at or below the contrast threshold of the cell in question. This was often the case with P cells where the SNR was << 1 at these contrasts, and a response average could not be determined that was distinguishable from noise. Such noise is not separable, and its inclusion in our SVD calculation might lead to the spurious conclusion that a neuron’s response is non-separable. To address this problem, we evaluated model fit by computing the normalized correlation coefficient (Schoppe et al., 2016) between the model fit and data with a criterion of 0.30 as a cutoff. This cutoff agreed best with visual inspection of firing rate traces and the instances where a condition did not appear different from Gaussian noise. We excluded conditions for which a cell’s fit fell below this criterion value.

Additionally, we excluded cells that did not pass this criterion at all luminance conditions and at least two contrasts. Altogether 20/75 cells did not meet these criteria, and the vast majority of these were parvocellular cells (18/20). The separability analysis is therefore based on the responses of the remaining 55 cells.

### Separable model evaluation

To evaluate the quality of the separable model fit to the neural data, we calculated the mean squared error between the separable and non-separable model predictions from the raw data. The non-separable model is simply the consequence of independently fitting an LN model to each condition. The separable model is derived from these model fits by singular value decomposition described above. We evaluated other methods to compare model predictions, including explainable variance and the normalized correlation coefficient (Schoppe et al., 2016). All methods gave very similar results.

## Results

We recorded neurons across layers of the LGN in 3 male anesthetized macaque monkeys using single electrodes. We assigned neurons to magnocellular layers (M layers) and parvocellular layers (P layers) based on eye preference and histological reconstruction, considering microlesions placed along recording tracks and depth. In certain instances, cells were encountered at boundaries between layers or within the S layers beneath magnocellular layer 1 (Kaas et al., 1978; Weber et al., 1983). These cells could have contrast gain signatures resembling P or M layer responses. However, the depth, eye preference, or chromatic signature (e.g., blue-yellow opponency) were consistent with the functional characteristics of koniocellular cells in the intercalated layers, as reported by others (Hendry and Reid, 2000). Therefore, we refer to them as putative koniocellular cells and denote them with **K\*** below. We only recorded a small number of these cells (n=12), and they are not the focus of this report.

### Contrast responses of M and P cells

We recorded 175 cells across layers in the LGN and performed an initial characterization of each cell using drifting gratings centered on the receptive field of each cell encountered. At each cell’s preferred spatial and temporal frequency, the response gain of M cells was far greater than that of P cells, as expected for cells in the M and P LGN layers (Derrington and Lennie, 1984; Kaplan and Shapley, 1982; Levitt et al., 2001; Movshon et al., 2005; Shapley et al., 1981) as well as M and P ganglion cells (Kaplan and Shapley, 1986; Lee et al., 1990).

We measured the contrast response function of each cell using drifting sinusoidal gratings of near-optimal spatial and temporal frequency, whose contrast varied in logarithmic steps from 0.01 to 1. The open circles in Figures **1A** and **B** illustrate the F1 contrast response function for typical M and P cells. Prior studies suggest three differences in contrast response distinguish M and P cell populations: the contrast level at which a cell reaches half its maximal firing rate (C_50_; vertical dashed lines in Figs. **1A** **and 1B**), the slope of the initial rise of the contrast response function (response gain), and the degree to which response saturates as contrast increases. We fit a descriptive model to the F1 response for each cell in our population. The model is a variant of the log contrast response function introduced by Robson (1975). It has the following form:

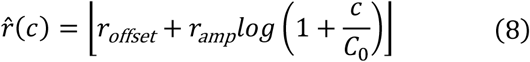

where *r*_offset_ <= 0 allows for a contrast threshold, *r*_amp_ > 0 is a gain term, *C*_0_ is a saturation constant representing the contrast at which logarithmic saturation begins, and *c* is contrast. Rectification prevents the function from having negative values. Excluding the offset term (*r*_offset_) led to underestimating the response gain and the C_50_ derived from this smooth function, particularly for cells that did not show responses at low contrast levels. We defined *response gain* as the slope from zero to the maximum contrast (*C*_sat_) at which the response is still linear. *C*_sat_ is the maximum contrast tested for linear cells, but it is the saturation constant (*C*_0_) for cells that saturate. We computed a saturation index from the raw data as follows:

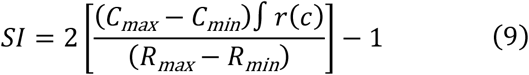

where *R*_max_/_min_ are the maximum and minimum response, *C*_max/min_ are the maximum and minimum contrast presented, and ⎰*r*(c) is the numerical integral of the contrast response function evaluated via the trapezoid method, as illustrated in Figure **1C**. Figure **1D** illustrates how this saturation index quantifies contrast response functions that are accelerating, linear, or saturating.

**Figure 1:**
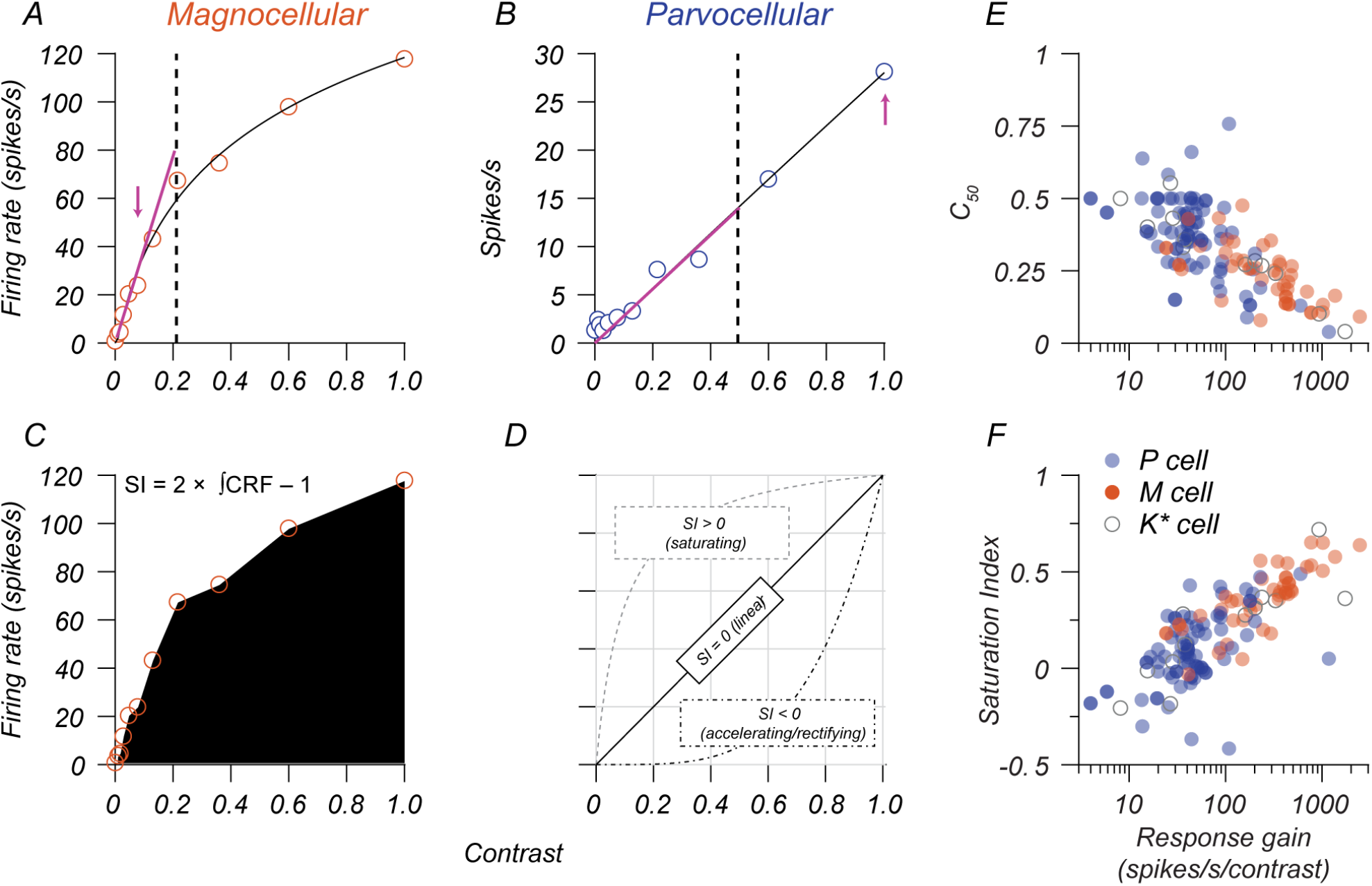
Quantifying response gain signatures of LGN neurons. **A, B**: Contrast response functions of an example M cell (A) and P cell (B). Open points are the first harmonic (F1) response at the temporal frequency of the stimulus. Smooth black lines indicate the fit of a descriptive function (equation 8) to these data. Dashed lines indicate the C50 (0.21 for M cell, 0.50 for P cell), and the magenta arrows indicate the maximum contrast within the linear range of each cell at which response gain was calculated (387 spikes/s/contrast for the M cell, 28 spikes/s/contrast for the P cell). **C**: Illustration of the method used to calculate a saturation index. The example M cell has a saturation index of 0.38, while the example P cell has a saturation index of -0.04. **D**: the relationship between the nature of the contrast response function (accelerating, linear, saturating) and the saturation index. **E, F**: Response gain vs. the C50 (E) and saturation index (F). Blue points represent P cells, red points represent M cells, and points with a gray outline represent putative K cells (K*).

Response gain and SI differed between P and M cell populations, with M cells having higher average contrast gain and more substantial saturation (t-tests, p<0.0001 for both). The median M cell response gain was 5 times the response gain of P cells, and the median C_50_ of M cells was 0.6x the C_50_ for P cells. Putative koniocellular cells (K*) had diverse response characteristics, with some responding like P cells and others responding like M cells. These results broadly match prior reports, except that several P cells in our population had a high response gain (>80 spikes/s/contrast). Further analysis shows that variations in the eccentricity of P cell receptive fields explain variations in their response gain, a factor we discuss below.

### How contrast and luminance influence responses in the retina

Previous reports have shown that both luminance and contrast gain control maximize sensitivity at high temporal frequencies. Increasing luminance selectively boosts contrast gain at high temporal frequencies in both M and P cells (Lee et al., 1990; Purpura et al., 1990). M ganglion cells exhibit a nonlinear dependence of contrast gain as a function of temporal frequency so that their contrast response function strongly saturates at low temporal frequencies and is linear at high temporal frequencies (Benardete and Kaplan, 1999). P ganglion cells have linear contrast response functions at all temporal frequencies (Benardete and Kaplan, 1997), so increasing contrast by a factor increases the firing rate by the same factor. We, therefore, expect the following for the contrast response functions of M and P-layer LGN neurons if they follow their retinal inputs.

1. M cell contrast response functions should saturate at low temporal frequencies and approach linearity at high temporal frequencies.
2. P cell contrast response functions should be linear at all temporal frequencies.
3. The response gain of M and P cells should increase with luminance, most prominently at high temporal frequencies,

### How contrast and luminance influence responses in the LGN

To evaluate whether our population of M and P cells were consistent with prior reports of retinogeniculate inputs, we recorded the response of M and P cells in response to drifting gratings that varied in contrast, luminance, and temporal frequency. We studied 75 cells (41 P, 24 M, and 10 K*) with the sweep stimuli (see Methods), in which temporal frequency varied over stimulus presentation time, as illustrated by the topmost curves in Figures **2A** and **2C**. The firing rates of two example cells are shown in gray below the stimulus profiles. We fit an LN model to these data and used it to estimate each cell’s amplitude and phase response, as illustrated in Figures **2B** and **2D**.

**Figure 2:**
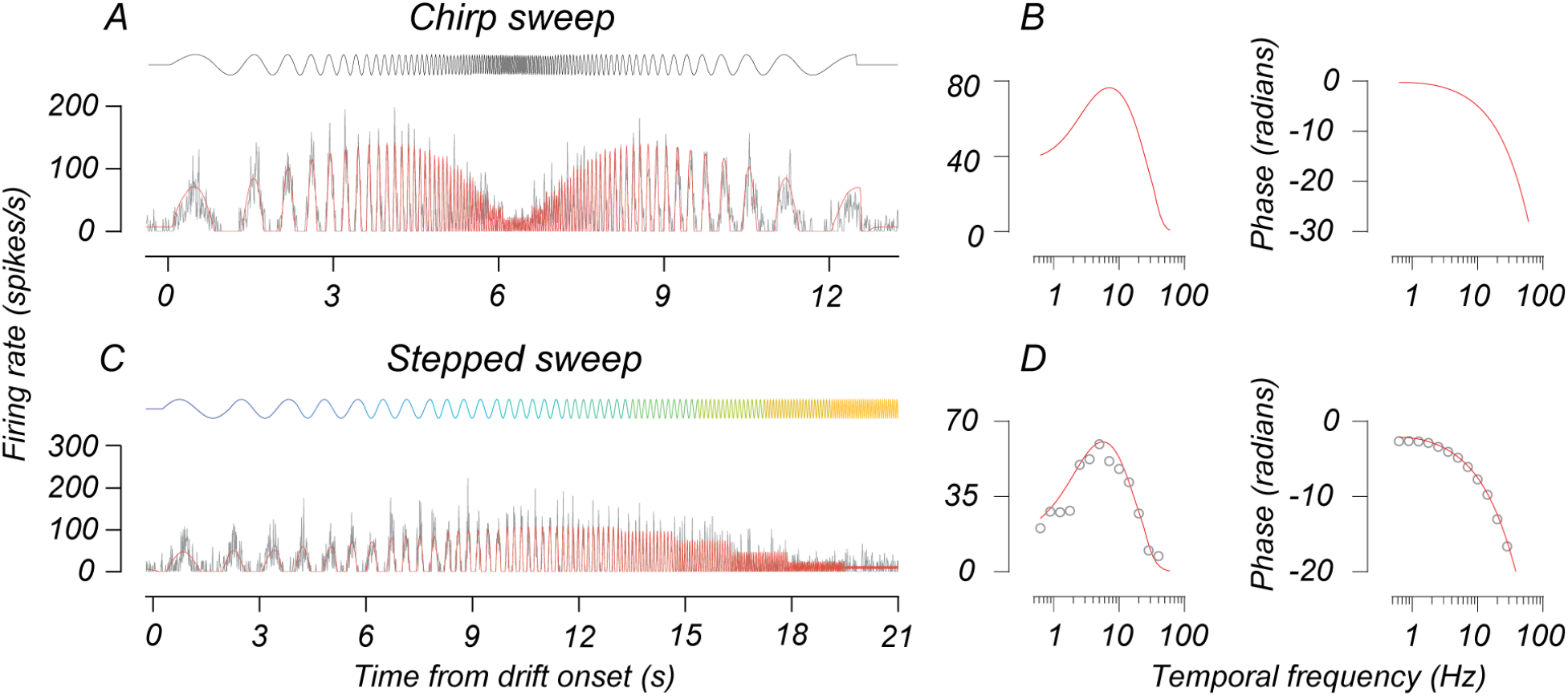
Using sweep stimuli to estimate temporal frequency tuning. **A, B:** The response of an example M cell to the chirp sweep stimulus. At the top of the figure, the black curve indicates the temporal contrast profile of the stimulus. In **A**, gray curves plot the cell’s firing rate over time, aligned to drift onset, and red curves are the response predicted by the LN model. The two plots in **B** give this LN model’s amplitude (left) and phase (right) response as a function of temporal frequency. **C, D:** The response of a different M cell to the stepped sweep stimulus. Plots follow the same conventions as A and B. The topmost multicolored sinusoidal curve in **A** indicates the temporal contrast profile of the stimulus. Colors indicate periods of constant frequency (each 1.6 s). The open red points in **D** are the amplitude and phase of the first harmonic response calculated from these spike times.

Figure **3** illustrates the temporal frequency tuning of two cells (one M and one P) at 3 luminances (from top to bottom) and 3 contrasts (from left to right).Open points illustrate the F1 response measured from these cells, and the smooth lines indicate the estimated F1 based on equation **1**. We averaged temporal frequency tuning curves across all conditions for each cell, selected the peak of this curve, and defined it as the “medium” temporal frequency. Frequencies 1.5 octaves below and above the medium temporal frequency were defined as “low” and “high,” respectively.

**Figure 3:**
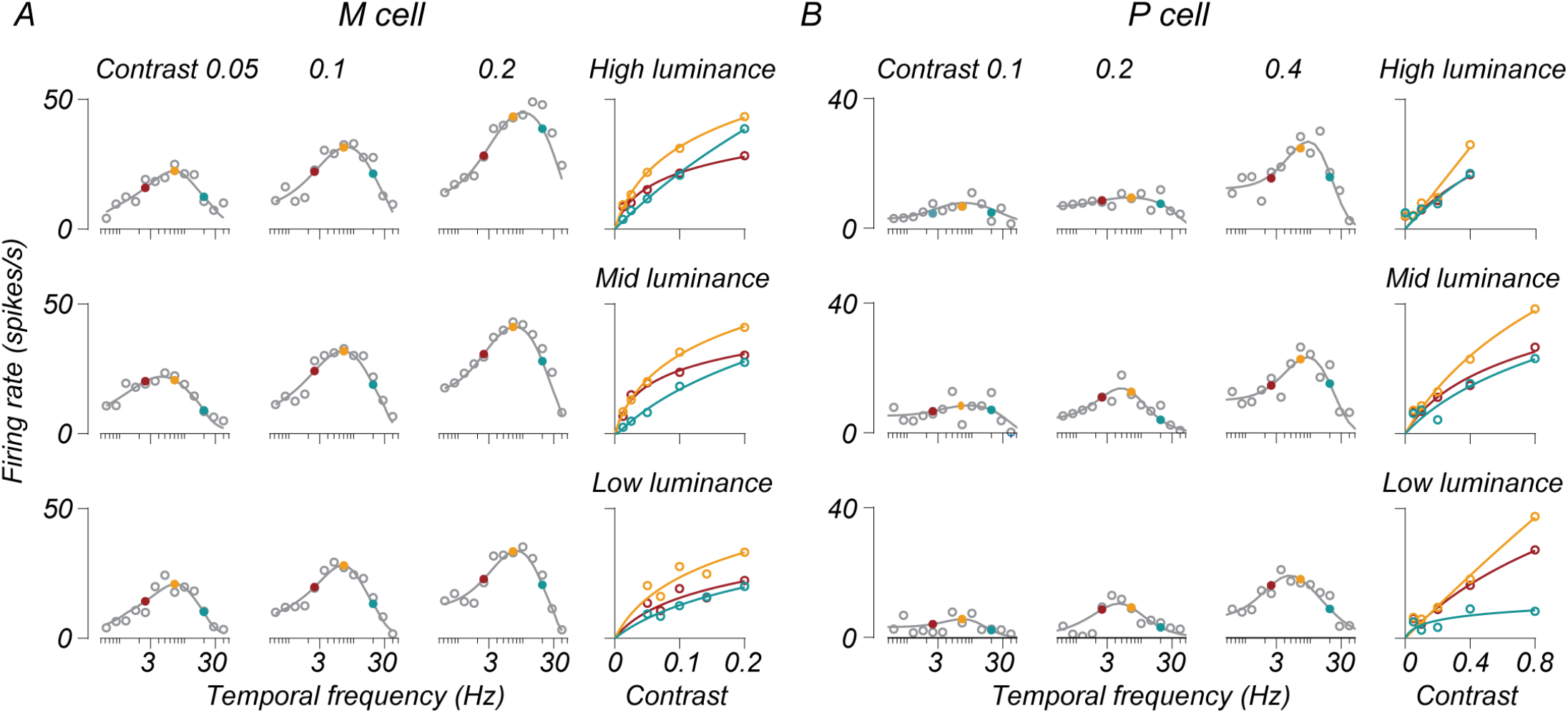
M and P cell temporal frequency tuning as a function of both luminance and contrast. **A, B**: The responses of one M cell and one P cell at multiple combinations of luminances and contrasts. In the main 3×3 plot, each subplot is a single set of measurements of temporal frequency tuning at a given contrast and luminance. Contrast values increase left to right, and luminance values increase from bottom to top. Open gray points are measured F1 responses for cells (recorded with the stepped sweep stimulus). Smooth lines through these points plot the amplitude response of the LN model fit to each condition. Filled points indicate temporal frequencies 0 +/- 1.5 octaves from the preferred frequency. Colors indicate low (red), medium (gold), and high (teal) temporal frequencies. The rightmost subplots in **A** and **B** show the contrast response functions estimated from the LN model at these three temporal frequencies (same color convention) for each luminance. Open points in these subplots are the LN model’s response as a function of contrast. The smooth curves in these subplots show the fit of a descriptive function (equation 8) to these data. Temporal frequency tuning curves in the main 3×3 plot show only a subset of stimulus contrasts, while the contrast response functions in the subplots indicate the response across all tested contrasts.

The filled red, gold, and teal points in Figures **3A** and **3B** indicate these temporal frequencies, for which we plot the contrast response functions. These two cells behave consistently with the observations of Benardete and Kaplan (1999). The M cell in Figure **3A** exhibits a saturating contrast response function for low and medium temporal frequencies but is more linear for high temporal frequencies (compare the red and gold curves to the teal ones in the rightmost subplots). The P cell in Figure **3B** has a more linear contrast response function across the evaluated temporal frequencies. Additionally, both cells show an elevation in response gain for high temporal frequencies as a function of luminance, consistent with (Purpura et al., 1990).

Across our recorded population, contrast response functions were broadly similar to these examples, but there was diversity across conditions. Figure **4** plots contrast response functions for all recorded M and P cells at low, medium, and high temporal frequencies at the middle luminance (10-12 cd/m^2^). Both M- and P-layer cells exhibited a wide range of response gains and degrees of saturation. The thick blue and red curves in Figure **4** are the averages of these functions. The population-averaged contrast response functions exhibit two clear trends. First, M cells had a higher average response gain and degree of saturation than P cells. Second, the difference between M and P cells was frequency-dependent: at high temporal frequencies, the contrast response functions of M cells are more linear (compare the initial slope and degree of saturation of the curves in Figures **4A** and **4B** with the curve in Figure **4C**). Separating population-averaged data by luminance condition revealed that the increased linearity exhibited by M cells at high temporal frequencies is luminance dependent.

**Figure 4:**
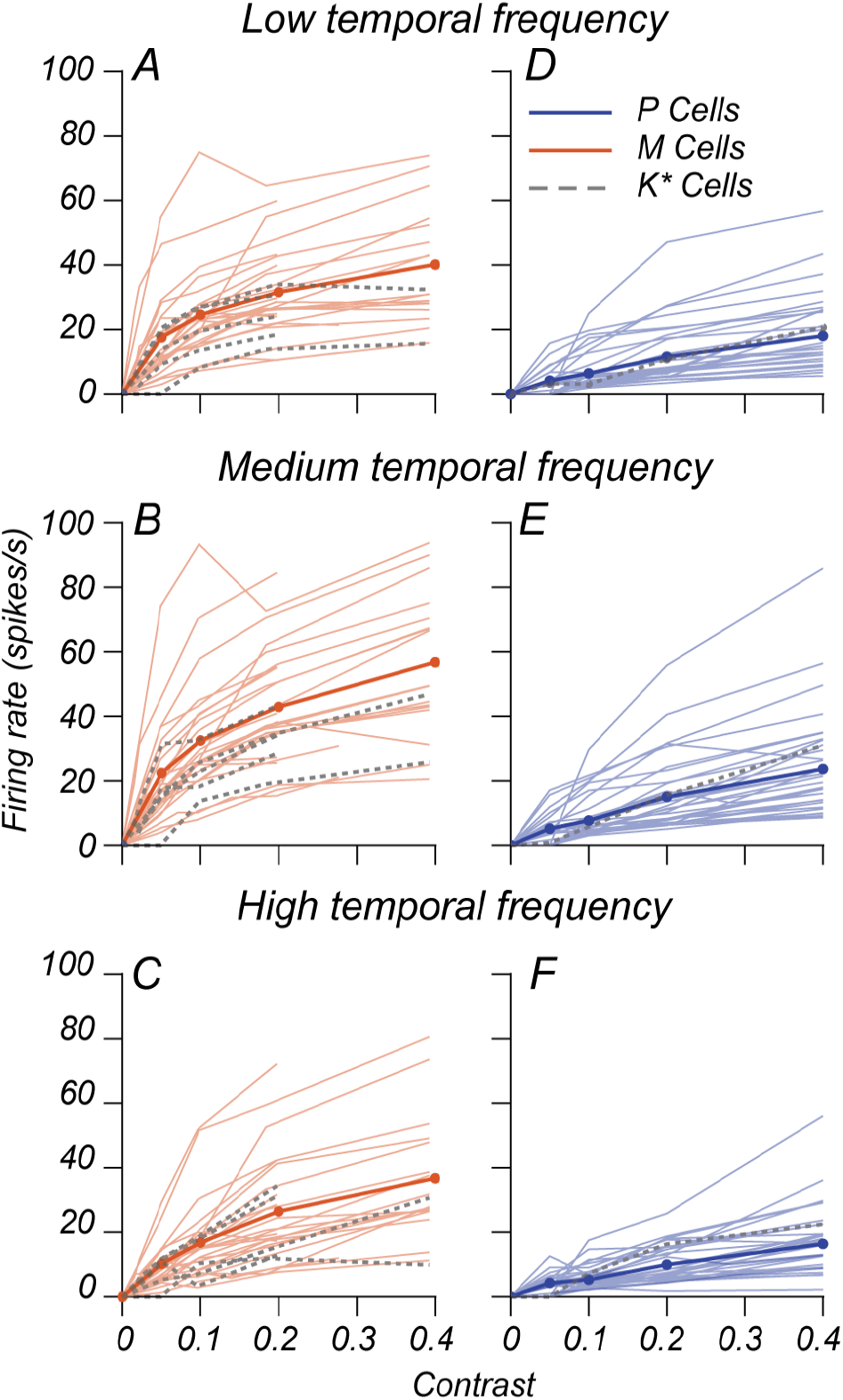
Diversity of contrast response functions within M and P cell populations. **A-F:** Contrast response functions from M and P cells recorded in the main luminance x contrast experiment. The thin lines in each subplot (**A-F**) represent data for one cell evaluated at contrasts between 0 and 0.4 using the LN model at one of three temporal frequencies indicated by the title at the top of each subplot. Blue lines represent P cells, red lines represent M cells, and dashed gray lines represent putative K cells. Thick blue and red lines indicate the population-averaged contrast response function. These data are taken from the mid luminance condition (10-12 cd/m2, 282-356 td). Given an initial characterization of each cell, the tested contrast ranges in the main experiment differed from cell to cell. Therefore, some M cell curves terminate at a contrast of 0.2.

The smooth lines in Figures **5A-F** indicate the population-averaged contrast response functions of M and P cells at different luminances. Individual curves in each subplot reflect the contrast response function evaluated at a particular temporal frequency (red = low, gold = medium, teal = high, as in Figure **3**). For both M and P cells, the contrast response function at high temporal frequencies (teal curves in Figures **5A** and **5D**) peaks at a low firing rate when stimulus contrast is 0.4, but as luminance increases (from left to right), the teal curves begin to lift upwards. For M cells, the consequence is that the contrast response function at low luminance has the same degree of saturation across temporal frequencies, while the contrast response function at medium and high temporal frequencies becomes increasingly linear and saturates less as luminance increases. P cell responses exhibit slight variation in saturation across changes in luminance but exhibit an increase in firing rate at 0.4 contrast as luminance increases. We found that this was not an effect of outliers. The bowtie plots in Figures **6A-F** depict the single-cell saturation indices of contrast response functions for low and high temporal frequencies relative to those for medium temporal frequencies. P cells (Figures **6A-C**) showed no systematic change in saturation due to frequency tested or stimulus luminance. By contrast, M cells (Figures **6D-F**) exhibit a systematic reduction in saturation index for high temporal frequency stimuli (*p* < 0.005, Wilcoxon signed-rank test), most prominently at medium and high luminances.

**Figure 5:**
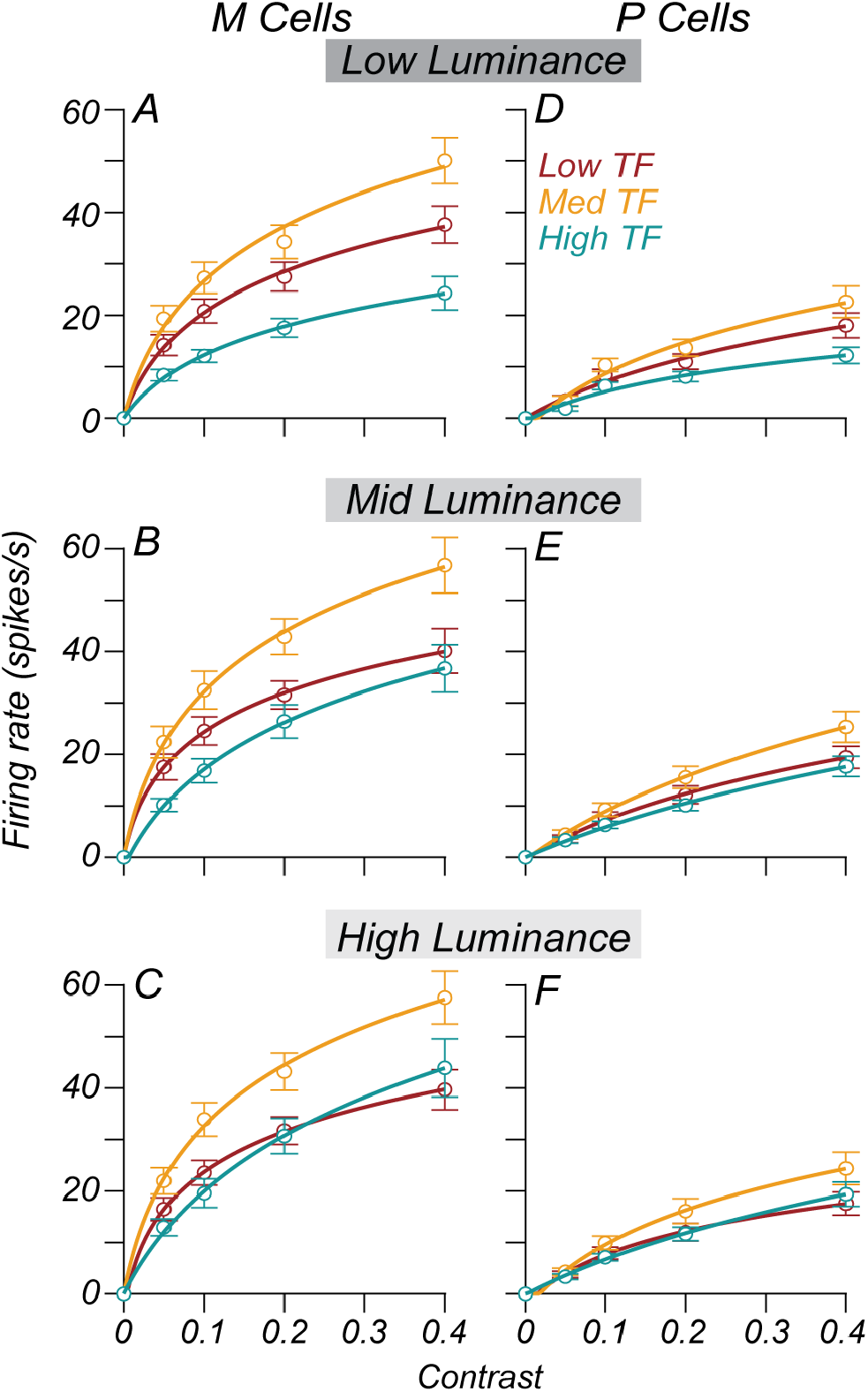
Average contrast response functions as a function of luminance. **A-F**: The average contrast response function for P cells (**A-C**) and M cells (**D-F**) over a contrast range of 0 to 0.4. The background luminance varies across rows. Low, mid, and high luminance correspond to 3.5, 12.6, and 41 cd/m^2^ for most cells (100, 356, 1159 td). The smooth curves through the points give the fit of equation 1 to the data. Error bars are the standard error of the mean across cells. Each curve’s color indicates the temporal frequency (TF). Gold = Medium TF, Red = Medium TF – 1.5 octaves, Teal= Medium TF + 1.5 octaves.

**Figure 6:**
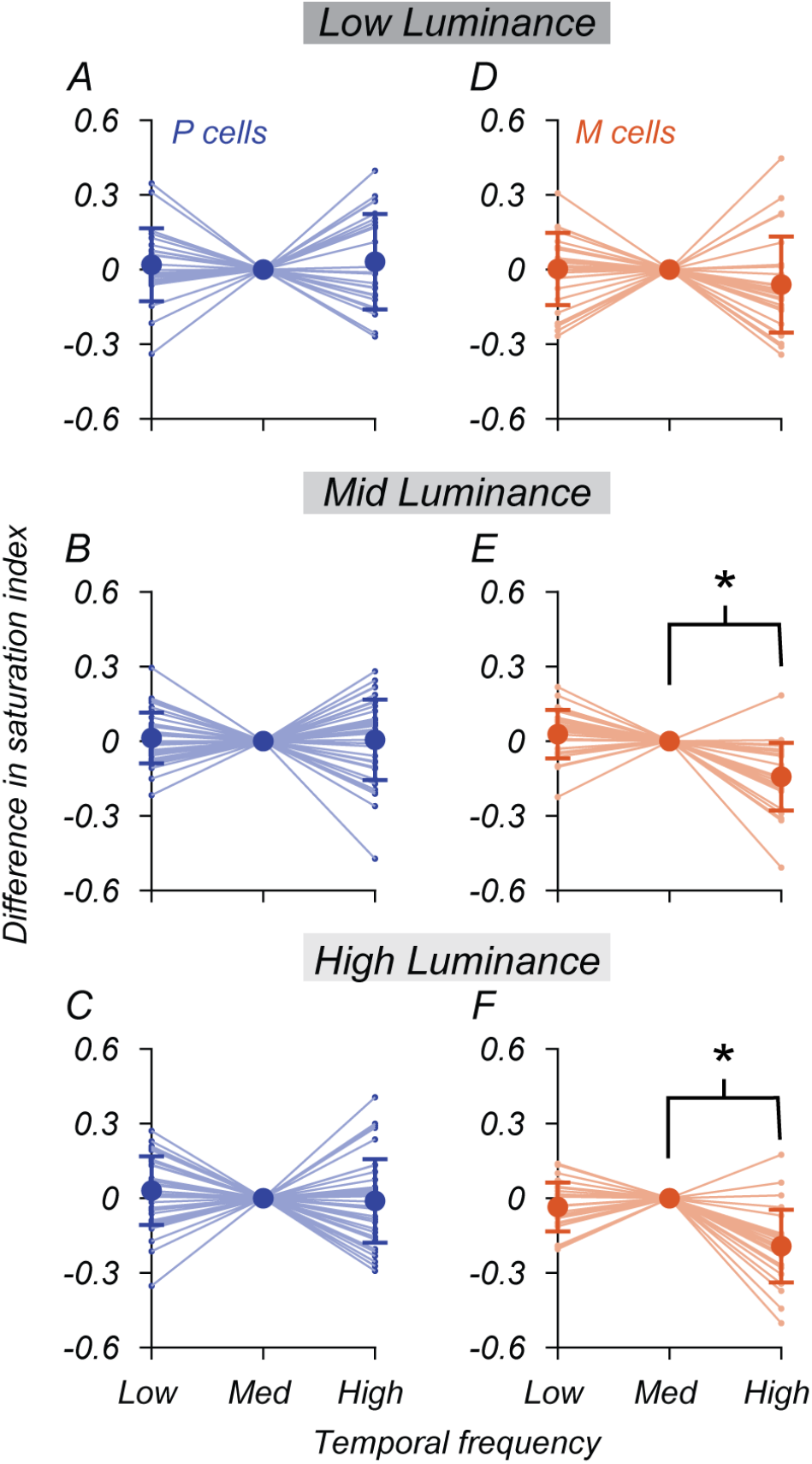
Change in response saturation as a function of luminance and temporal frequency. **A-F**: Each thin blue or red line in these figures indicates one cell’s saturation index calculated at one of three temporal frequencies. Saturation indices for low and high TF stimuli are displayed relative to the saturation index for medium TF stimuli. Each row corresponds to a different luminance level, increasing from top to bottom as in Figure 5. Blue lines represent P cells, and red lines represent M cells. Filled points indicate the mean saturation index per temporal frequency condition, and the error bars on each point indicate the standard error of the mean. Asterisks indicate comparisons between conditions (indicated by brackets) for which a Wilcoxon rank-sum test had a p-value < 0.005.

Both M and P cells’ contrast response functions tend to reach a higher firing rate when both luminance and temporal frequency are high. This seems consistent with the expectations based on retinal data. Estimates of the slope of the contrast response function proved highly variable, however.

In our dataset M and P cells tend to saturate at low firing rates for low luminance stimuli (see figure 5), and the confluence of limited contrast conditions and measurement noise made it difficult to constrain model parameters during optimization. We, therefore, analyzed the influence of luminance on response gain by considering the firing rate at 0.2 contrast for each cell. This contrast is within the linear range of nearly all the M and P cells we recorded, so if luminance causes an increase in response gain, the firing rate at this contrast should increase as luminance increases. The bowtie plots in Figure **7A-F** illustrate the change in firing rate at 0.2 contrast for individual cells in low and high luminance conditions relative to the medium luminance condition. There was almost always a consistent increase in firing rate at 0.2 contrast for both M and P cells as luminance increased from low to medium levels (3.5 to 12 cd/m^2^). For medium and high temporal frequency stimuli, the average firing rate for M cells at 0.2 contrast increased 8.73 spikes/s (p < 0.005, Wilcoxon rank-sum test) as luminance increased from the lowest to mid-levels. The same was true for P cells, but the firing rate increased by 3.95 spikes/s (p < 0.005, Wilcoxon rank-sum test) as luminance went from the low to the middle level.

**Figure 7:**
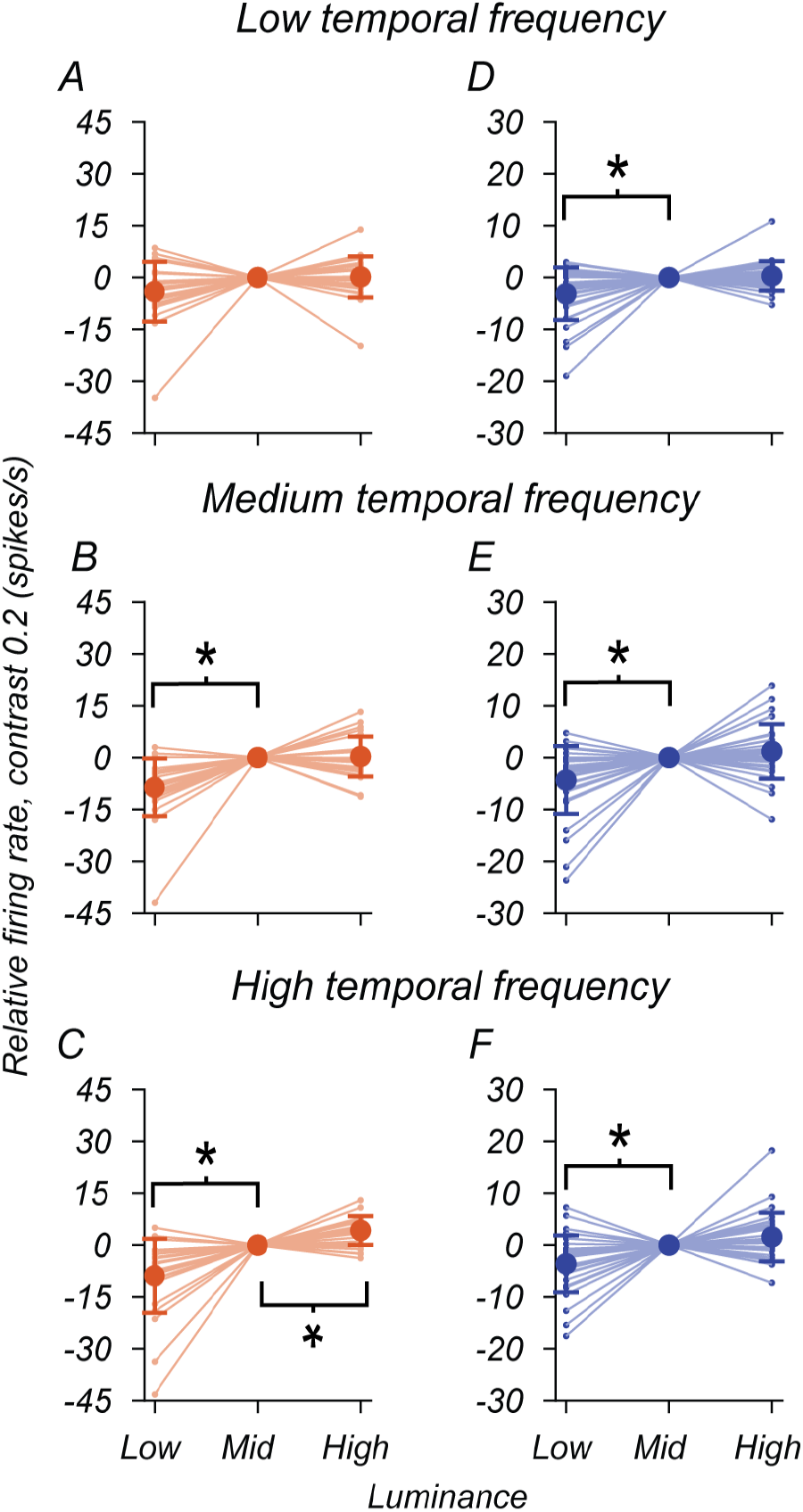
Change in firing rate as a function of luminance and temporal frequency. **A-F:** Firing rate at 0.2 contrast as a function of mean luminance and temporal frequency. Each subplot shows the firing rate at a contrast of 0.2, for one temporal frequency, and one cell type, estimated from the LN model and a fit of equation (5). Each row corresponds to a different temporal frequency, increasing from top to bottom. Each thin line is one cell’s response gain across three luminance levels. Blue lines represent P cells, and red lines represent M cells. The firing rates for high and low luminance conditions are shown relative to the firing rate in the mid-luminance condition. Asterisks indicate comparisons between conditions (indicated by brackets) for which a Wilcoxon rank-sum test had a p-value < 0.005.

Together these results suggest that cells in the LGN behave similarly to retinal ganglion cells. The contrast response function for high frequency and low luminance stimuli saturates early, plateauing at a low firing rate (see M and P cell responses in Figures **3A** and **3B**). The firing rate analysis in Figure **7** is consistent with the results of Purpura et al. (1990). They reported that responsivity, the firing rate of a cell at the peak of its linear range divided by the contrast, increased as a function of luminance. While the results of Lee et al. (1990) suggest that this is a consequence of an increase in the initial slope of the contrast function as a function of luminance at high frequencies, we could not confirm this result, given our limited measurement time.

### Separable influences of contrast and luminance on temporal frequency tuning

The prior results illustrate how M and P cells’ response gain and saturation change as a function of temporal frequency and mean luminance. The similarity of these results with prior reports of the retinal input to the LGN (Benardete et al., 1992; Lee et al., 1990) suggests that these adaptation mechanisms are retinal in origin. In the retina, the mechanisms responsible for adapting retinal ganglion cells to a given mean light level appear mostly upstream of those believed to define modulation sensitivity at different contrasts (Shapley and Enroth-Cugell, 1984).

If the adaptation observed in LGN neurons were a consequence of adaptation in retinal inputs, it would predict that mean luminance and contrast changes will have relatively independent effects on the amplitude of response at a given temporal frequency for LGN neurons. Mante et al. (2005) tested this explicitly and concluded that in cat LGN, the effects of luminance and contrast on the responses of LGN cells are independent. We extended this approach to M and P populations in the monkey and found that we could decompose temporal frequency tuning into a separable set of factors that vary with luminance or contrast alone.

Figures **8A** and **8B** depict the tuning of the same M and P cells initially described in Figure **3**. The smooth purple lines running through the data points (open gray circles) are the product of two sets of tuning curves, one varying by luminance alone and the other by contrast alone. These separable tuning curves appear under shaded blue and red boxes above the points. We estimated these separable functions using singular value decomposition (SVD), as done in the study of Mante et al. (2005). Our analysis begins with a matrix *F*_*L,c*_ (*ω*) that describes the first harmonic response (in the frequency domain) observed at a given temporal frequency (*ω*), contrast (*C*), and luminance(*L*). For example, consider the matrix containing the tuning curve data presented in Figure **3**. By applying SVD to his matrix, we factorize it into two separable matrices, *F*_*C*_ (*ω*) and *F*_*L*_ (*ω*). Each of these matrices will represent a triplet of temporal frequency tuning curves that vary as a function of either contrast or luminance and whose product is the best separable approximation to the data. We refer to this decomposition as the separable LN model henceforth. For the two cells shown in Figures **8A** and **8B**, the separable LN model generated using SVD accounted for 94% of the variance in the data.

**Figure 8:**
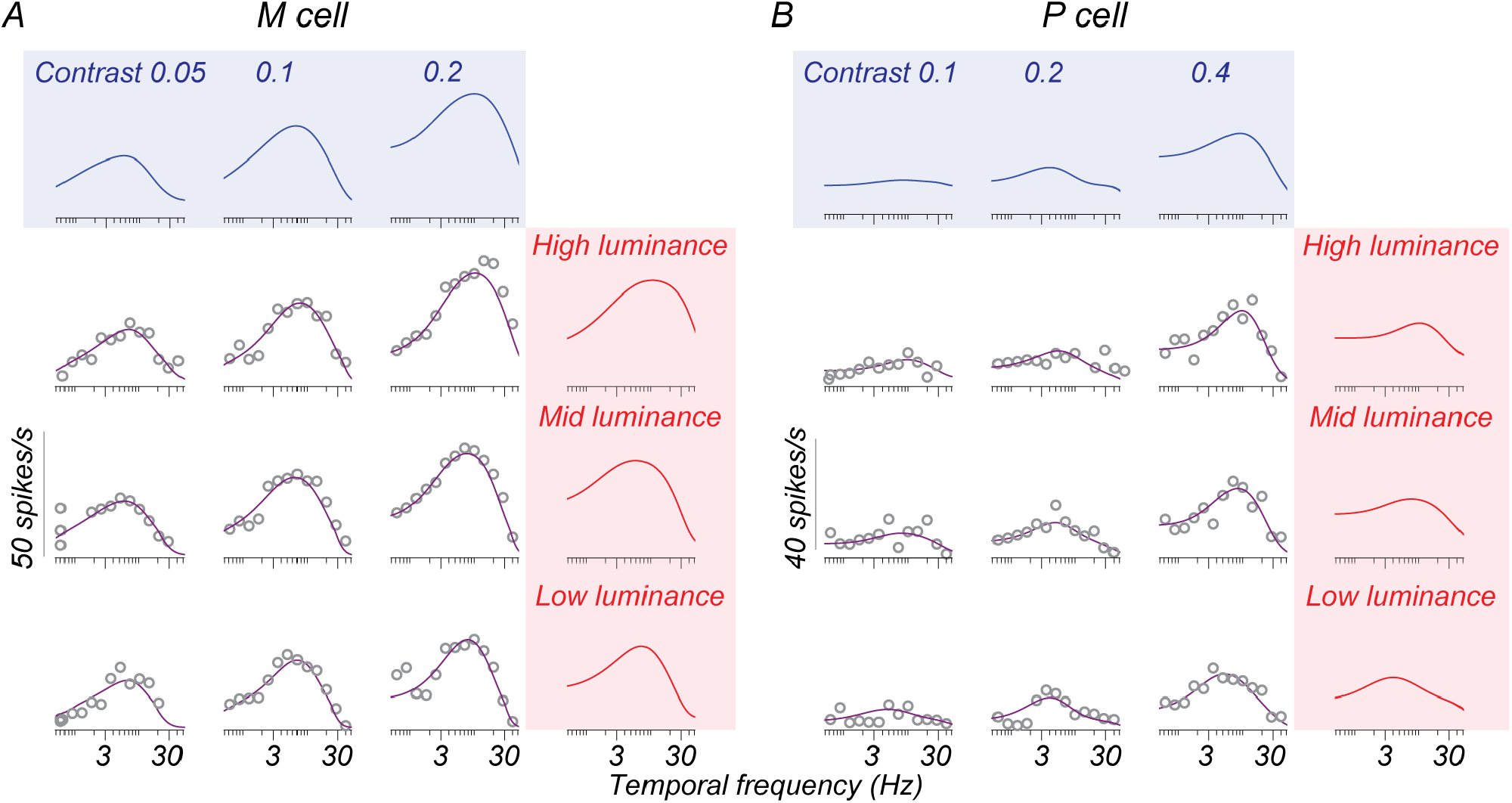
Decomposition of temporal frequency tuning into separable effects of luminance and contrast. **A, B**: Temporal frequency tuning of example M and P cells for the stepped sweep stimulus and the separable LN model prediction. Gray open points in the area bounded by the shaded boxes are the F1 response at different temporal frequencies recorded from each cell at a given luminance (row) and contrast (column) condition. These data are those appearing in Figure **3**. The solid purple line is the separable LN model’s prediction of the F1 response in each condition. The solid blue and red curves in the shaded boxes above these tuning curves are separable functions of contrast and luminance derived using singular value decomposition. The outer product of the separable functions gives the purple lines.

As illustrated in Figure **2**, the F1 response in the frequency domain is directly related to a cell’s response in the time domain. By performing SVD in the frequency domain and inverting the separable predictions into the time domain, we could generate a prediction of the average firing rate over time in any condition (Figure **9**). In Figures **9A** and **9B**, the gray curves illustrate the average firing rate over the stimulus presentation period of one condition (41 cd/m^2^, 0.2 contrast) for the M cell introduced in Figures **3A** and **8A**. The orange curve in Figure **9A** shows the fit of the original LN model to these data (before the application of SVD). In Figure **9B**, the purple curve shows the firing rate prediction generated using the separable LN model.Comparing data in the time domain allowed us to collapse across the datasets where we could compute the F1 directly from the data (the stepped sweep data) and the datasets we could not (the chirp sweep data). The original and separable models do an equivalent job capturing response for this cell such that the root-mean-squared error (rMSE) between the data and each model’s prediction is very similar (15.35 spikes/s for the original model vs. 15.45 spikes/s for the separable model). Across the population of recorded cells, the performance of the separable model was essentially indistinguishable from the performance of the original LN model. Consequently, nearly all points lie along the unity line when we plot the rMSE for the original LN model vs. the rMSE for the separable model (Figure **9C**).

**Figure 9:**
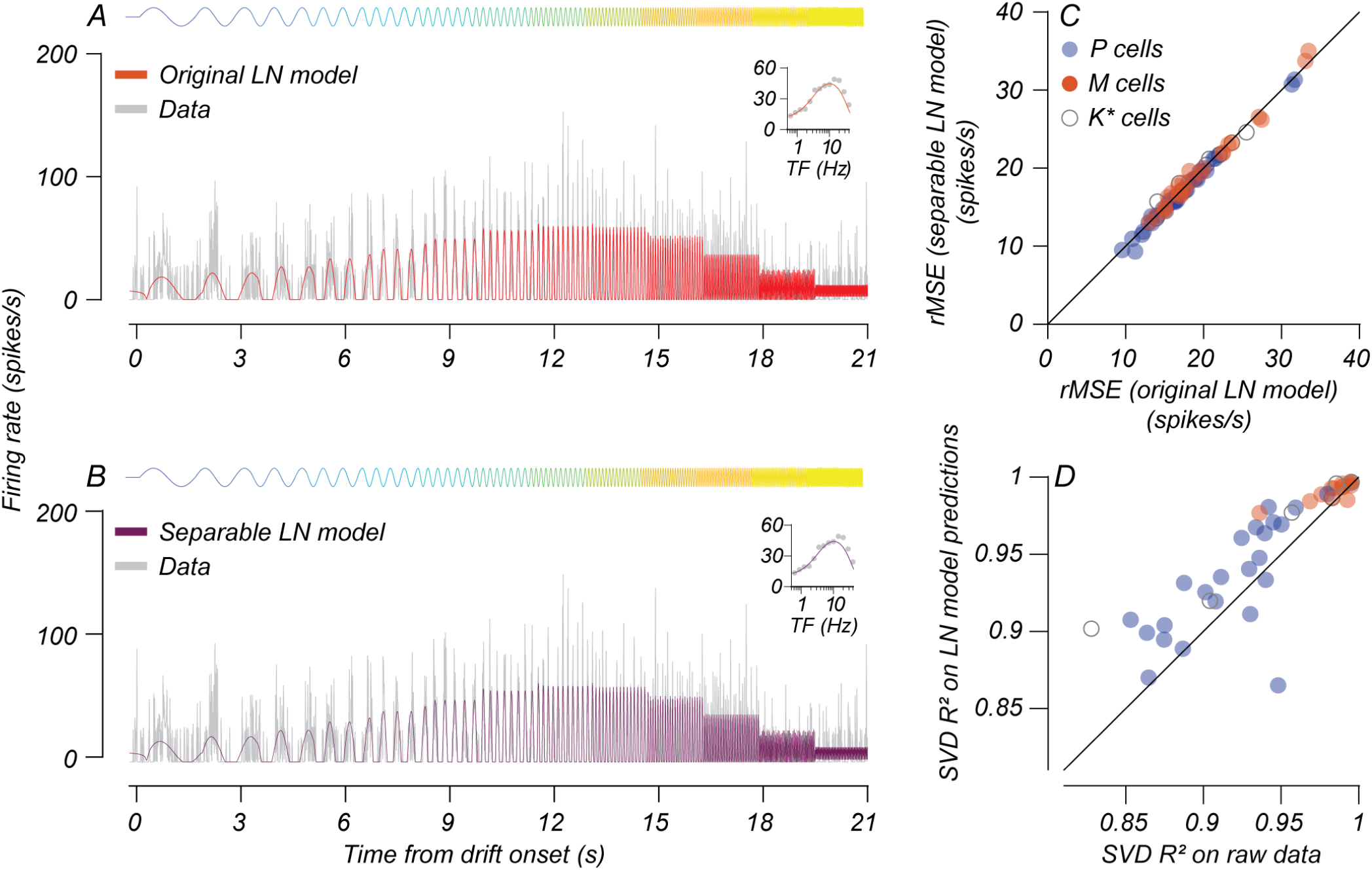
Comparing separable model predictions vs. original LN model and F1 responses. **A, B:** Original and separable LN model predictions for an M cell recorded during a stepped sweep in one stimulus condition. The gray curves indicate the cell’s firing rate over time and are the same in both plots. The curve above each figure indicates the change in stimulus contrast in the receptive field over time. The solid orange and purple curves inset in **A** and **B** are the predictions of the original and separable LN models, respectively. The root-mean-squared errors (rMSE) between the original and separable models and the measured response were 15.35 and 15.45 spikes/s, respectively. **C:** The root-mean-squared error between the original (x-axis) or separable (y-axis) LN models and the cell’s firing rate. Each point indicates a single cell, and colors indicate cell type as in previous plots. The black line is unity. **D:** Variance explained (R^2^) by SVD on raw F1 response vs. variance explained (R^2^) on F1 responses predicted by the LN model. The data follow the same conventions used in **C**.

### Controls for model fit quality

One limitation of these analyses is that they rely on the fit of a model to generate the separable tuning curves. To see if this biased the results presented in Figure **10C**, we analyzed the responses of cells on which we ran a stepped sweep, gathered in 2/3 of our animals (n=42 cells). As stated above and in the methods section, the first harmonic response for this stimulus can be computed directly from spike times without relying on a model fit to the data. The F1 response at 13 tested temporal frequencies at multiple luminance and contrast conditions yields a matrix *F*1_*L,C*_ (*ω*), similar to the one described above. SVD on this matrix yields two separable matrices, *F*1_*C*_ (*ω*) and *F*1_*L*_ (*ω*), that capture tuning curve shifts driven by contrast or luminance alone and whose product provides the best separable approximation to the raw F1 data. We computed the variance accounted for (R^2^) by singular value decomposition applied to either raw F1 data or LN model predictions for those same temporal frequencies. Figure **9D** illustrates that the SVD accounts for an equal share of the response variance for either set of measurements. This analysis suggests that our LN model fitting procedure did not bias the SVD towards separable response functions. One caveat is that SVD on raw F1 had an average R-squared value of 86% across P cells and 95% across M cells (Figure **9D**). We do not think this is a material difference indicating that P cells are less separable. Rather, the lower R-squared values of P cells are a consequence of the greater variability/noise of P cell responses at lower contrasts that fell at or below these neurons’ contrast threshold.

**Figure 10:**
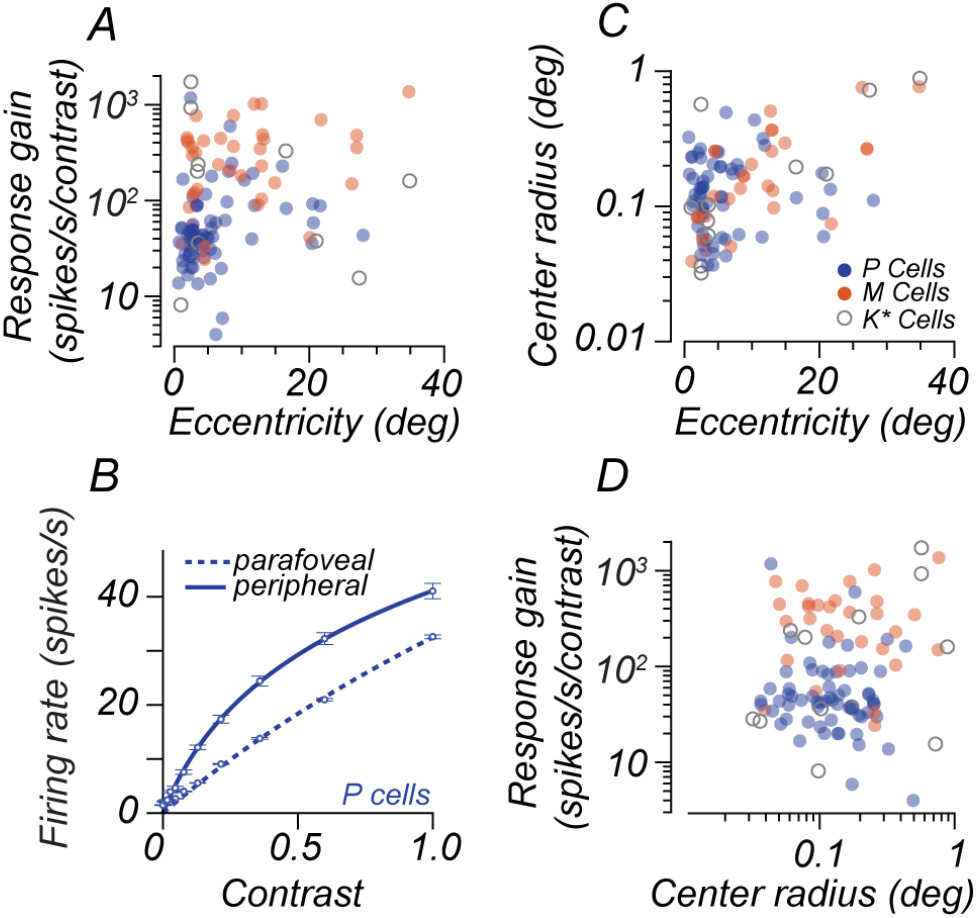
Variation in response gain with receptive field size and eccentricity. **A:** Response gain for each cell vs. receptive field eccentricity **B:** The population-averaged contrast response function for parafoveal vs. peripheral P cells. **C:** Center RF radius as a function of eccentricity. **D:** Response gain plotted as a function of center RF radius for each cell type. Each point in Figures **A, C**, and **D** is one cell. Blue points represent P cells, Red points represent M cells, and points with a gray outline represent putative K cells (K*). The data and solid curve in **B** show the population-averaged contrast response function across parafoveal P cells (eccentricity < 8), and the data and dashed curve show the population-averaged contrast response function for peripheral P cells (eccentricity > 8 degrees).

### Eccentricity accounts for diversity in P cell response gain

The previous analyses quantify how response gain varies as a function of temporal frequency and luminance in both cell classes. However, while they capture how these factors influence each cell class on average, they do not fully account for the diversity of response gain signatures within a given cell class. More specifically, it is unclear why P cells exhibit higher variation in response gain than M cells in our dataset (consider the P cell response gain range in Figures **1E** and **1F**). While most P cells in our population have response gain characteristics consistent with prior reports, namely a low response gain, there are several examples in which this is not the case. We found that the eccentricity of P cell receptive fields covaried with their response gain.

The LGN cells in this study had receptive fields spanning eccentricities ranging from 0.5 to 35 degrees from the fovea. Figure **10A** plots the response gain data presented in Figure **1** against receptive field eccentricity. Most cells we recorded were parafoveal (within the central 8 degrees of the fovea), spanning response gains from 5-200 spikes/s/contrast. Cells recorded further out in the periphery (beyond 8 degrees in eccentricity) had response gains closer to the right tail of response gain distribution in the parafovea. As a result, the population-average contrast response function for P cells recorded further in the periphery had a higher response gain and saturated more than the population-average contrast response function recorded in the parafovea (Figure **10B**). With the caveat that we only recorded a small sample of P cells beyond 8 degrees in eccentricity (n=11), there was a significant (Wilcoxon rank-sum, p < 0.005) difference between response gain in these two groups. M cells did not show such a difference in response gain as a function of eccentricity.

The dendritic field size of major ganglion cell classes projecting to the LGN increases with eccentricity (Dacey and Petersen, 1992). Consequently, the receptive field center size (RF center) also increases with eccentricity (Derrington and Lennie, 1984). Cells with larger RF centers absorb more light (the effective flux is higher over the center mechanism). In the cat, the effective flux determines the adaptation state and the response gain of the ganglion cell inputs to the LGN, such that cells with larger RF centers have higher response gain (Enroth-Cugell and Shapley, 1973; Shapley and Enroth-Cugell, 1984). Prior work has suggested that a similar mechanism might account for the response gain differences between M and P cell populations (Purpura et al., 1988); therefore, we wondered whether center RF center size would predict P cell response gain. While we confirmed prior reports (Derrington and Lennie, 1984) that M and P cells differ in center RF size (on average), we found no clear relationship between response gain and center mechanism size within either cell group. We estimated center RF size from the fit of a difference of Gaussians to the amplitude response of each cell to gratings of different spatial frequencies (Enroth-Cugell and Robson, 1966).

The scatter plot in Figure **10C** plots the center RF radius of every recorded cell against its receptive field eccentricity. The center area for M cells grew with eccentricity (*r* = 0.68, p < 0.005). There was no correlation between P cell center size (*r* = 0.01, ns) and eccentricity. Finally, center RF size and response gain were uncorrelated in M and P cells (r = 0.29 and -0.05, both ns; Figure **10D**).

## Discussion

Our results show that responses in the magnocellular and parvocellular layers of the non-human primate broadly follow the documented properties of retinal inputs to these layers. Luminance and contrast gain control make separable contributions to steady-state responses driven by sinusoidal stimuli. Increasing luminance leads to an elevation and linearization of response gain for high temporal frequency stimuli. The contrast gain control’s effect seems relatively independent of luminance, as saturation for stimuli at or below individual cells’ peak temporal frequency preference occurs regardless of the background light level. Our findings are consistent with the results of retinal recordings (Benardete et al., 1992; Benardete and Kaplan, 1999, 1997; Purpura et al., 1990). Separability of these two processes can be demonstrated directly on temporal frequency responses measured at different luminances and contrasts, extending the results reported by Mante et al. (2005) from the cat LGN to the primate LGN.

### Response gain and saturation for M and P cells at low luminance values

An unexpected finding illustrated in Figure **5** is that many M and P cells appear to plateau at a low firing rate for high temporal frequency stimuli under low luminance conditions. Firing rates saturate at low contrast levels, making response gain measurements more variable. The result is that cells respond less strongly to high temporal frequency stimuli at low luminance compared to high luminance. Prior studies have not reported this interaction between saturation and luminance for high-frequency stimuli. The work of Benardete and Kaplan (1999), which reported a linear contrast response for high TF stimuli, was carried out using similar retinal illuminances to our highest value. There are three possibilities as to why this is. The first two are methodological:

1. The lowest contrast we presented for many of our cells was 0.05. For cells whose responses saturate by 0.05, the initial slope of the contrast response function is poorly constrained and makes it difficult to obtain a reliable estimate of response gain.
2. The decreased responsivity at low luminance levels may make our estimates more vulnerable to noise. It is possible that with additional stimulus presentations, the contrast response function for high-frequency stimuli might appear more linear.
3. The saturation we observe may indicate an additional gain control mechanism endogenous to the LGN that saturates high temporal frequency responses.

We view this last point as the least likely, but our data at present are not sufficient to rule it out. Our experiments made a trade-off between the number of repeats and the number of conditions we surveyed. Given the results above, however, we could increase recording time and rule out response variability by using stimuli with a more restricted set of temporal frequencies and the same range of luminance and contrast conditions.

### The origin of eccentricity-dependent effects

For P cells, response gain tended to increase as we recorded at greater retinal eccentricities. As a result, outside the parafovea, the distribution of response gain for both M and P cell classes became more similar (Figures **1** **and 10**). Given that center receptive field size is reported to increase with eccentricity (Derrington and Lennie, 1984), one might predict a higher response gain simply as a consequence of greater effective luminous flux over the receptive field center (Enroth-Cugell and Shapley, 1973; Purpura et al., 1988; Shapley and Enroth-Cugell, 1984). From this perspective, our results are puzzling. First, we found no increase in P cell receptive field size as a function of eccentricity (see Fig. **10C**). Second, there was only a weak covariation between center area and response gain within each cell class. We compared our data to a previously published dataset in our lab (Movshon et al., 2005) to address these criticisms, suspecting that a sampling bias accounted for this problem.

The results in Figure **11** show M and P cells recorded in each dataset using different symbols (**x** and **o**). Most of the data presented in this manuscript (**o**) fell within the bounds of this previous dataset. Figure **11A** shows the relationship between eccentricity and response gain for the two datasets (226 cells total, 123 P cells, 91 M cells). M cell response gain did not vary as a function of eccentricity (Spearman’s r=0.16, ns). By contrast, P cell response gain did increase as a function of eccentricity. For cells for which we measured spatial frequency tuning (262 cells total, 145 P cells, 105 M cells), there was an increase in center radius with eccentricity (Figure **11B**). This increase was the case both for P cells (Spearman’s r = 0.45, p< 0.005) and M cells (Spearman’s r = 0.59, p< 0.005). A few outlying P cells in the current dataset had larger than expected center radii. These cells were all recorded near the fovea (within 1.5 degrees). We think these results are primarily due to measurement error stemming from imperfect correction of the animal’s optics. It is also possible that small eye movements under anesthesia biased results. Together, these factors and the results in Figure **11** are consistent with the observation that the near-foveal RF estimates are less reliable than estimates at or beyond 3 degrees from the fovea (Derrington and Lennie, 1984).

**Figure 11:**
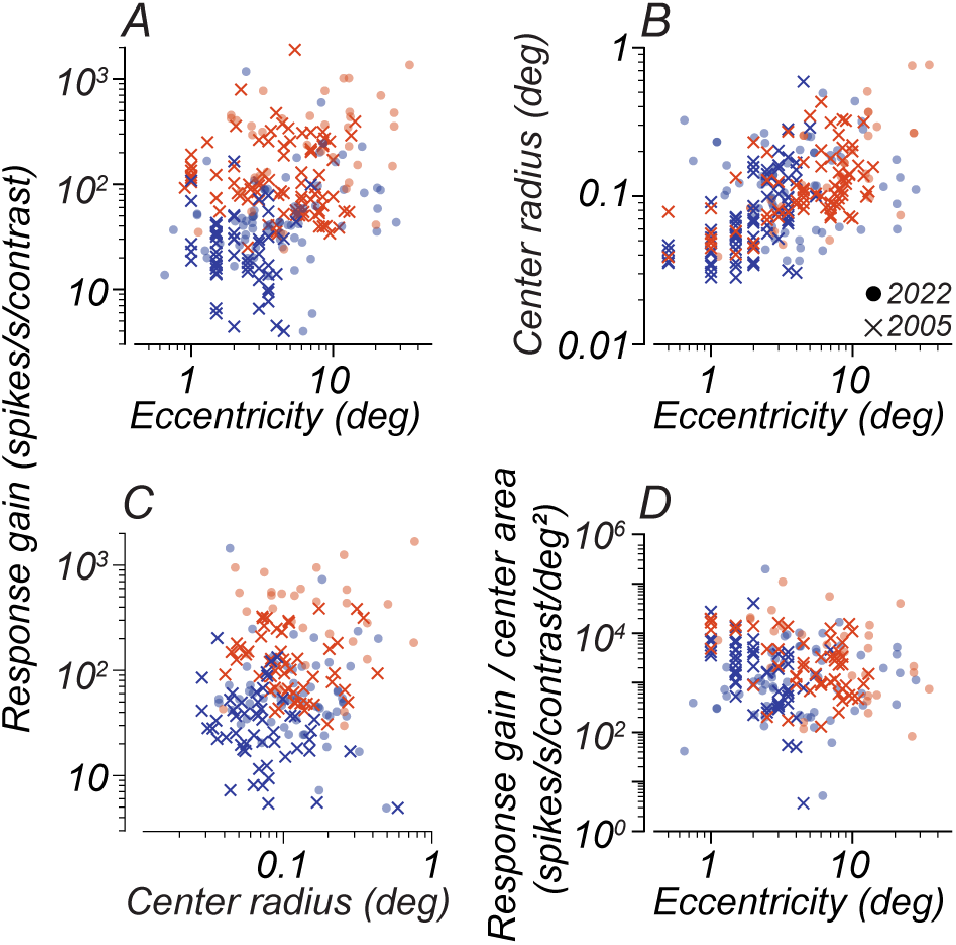
Response gain for two separate data sets from this laboratory. **A:** Response gain as a function of eccentricity **B:** Center radius as a function of eccentricity. **C:** Response gain plotted as a function of center radius and for each cell type. **D:** Normalized response gain (gain/center area) plotted as a function of eccentricity. Each point in figures **A-D** represents one cell and follows the conventions of Figure **10**. Blue points represent P cells, and red points represent M cells. Filled circles indicate cells recorded in the current study, and cross marks indicate data taken from the adult recordings reported by Movshon et al. (2005).

There was no covariation between receptive field radius and response gain for either M or P cell classes (Fig. **11C**) even with this larger dataset. We reconciled these findings by considering the response gain of cells normalized by their center area. We plot this normalized gain as a function of eccentricity in Figure **11D**. It is apparent that accounting for center size brings M and P cells into greater alignment across eccentricities (compare Figs. **11A** vs. **11D**). This alignment suggests that center size differences may be a mechanistic explanation for some differences between M and P cells. Additionally, we found that beyond 8 degrees, there was no difference between M and P cell normalized response gain (Wilcoxon rank-sum test, p = 0.9667). Interestingly, within 8 degrees, M cell normalized response gain was roughly 4-fold greater than P cells (Wilcoxon rank-sum test, p < 0.005).

In summary, at least part of the variation in response gain across eccentricity and cell class might be accounted for by the center receptive field size, as hypothesized initially by Purpura, Kaplan, and Shapley (1988). However, within the central 8 degrees, some additional factor contributes to M cells having higher response gain than P cells. Cone inputs and center-surround delays are known to change with eccentricity (Lee et al., 2012; Solomon et al., 2005), and these factors might account for the higher than expected response gain of M cells within the central 8 degrees. For small receptive fields nearer to the fovea, optical blur can confound. In future work, mapping cone inputs to receptive field centers and surrounds (Field et al., 2010; Reid and Shapley, 2002) could be carried out conjointly with estimates of response gain to explore this idea quantitatively.

An additional factor is the image blur caused by the eye’s optics that attenuates the response to high spatial frequencies (and corrupts estimates of center size by sinusoidal gratings). Consequently, P cell receptive fields near the fovea will have a bias towards larger estimated center receptive field sizes (Lee et al., 1998). M cells will be less affected by the eye’s optics due to their larger size. In theory, future studies can measure the point spread function of each animal’s eye and use tailored stimuli (McMahon et al., 2000) to bypass this optical blur and provide a more accurate estimate of center receptive field size.

### Gain control in other pathways

Throughout this report, we have included data from putative koniocellular cells. Some of these cells behave like our identified P layer cells, others like identified M layer cells. Concrete conclusions are difficult to draw, given the limited sample of recorded K cells. However, this leaves the opportunity for future work characterizing this population and others. Indeed, while most retinal input to the M- and P-layers of the LGN is from the midget and parasol ganglion cells in the retina, many other ganglion cell types project to the LGN from the retina (Dacey, 2004). Some of these terminate in the intercalated (koniocellular) layers, and some terminate in the P and M layers (Crook et al., 2008; Dacey, 1994), and each presumably has its own gain control signature – but there have yet to be studied. Isolating different cell classes is challenging, especially as we do not have functional signatures or laminar boundaries for many of them that allow their identification. However, there are two cell classes with relatively well-understood and distinct functional signatures whose gain control characteristics we can investigate. One class is the blue-yellow opponent cells that terminate within the koniocellular layers (Dacey, 2004). We only recorded two blue-yellow opponent cells in the data reported here, but experiments targeting just the blue-yellow population are possible. The second class is the smooth monostratified cells that project to the M layers (Crook et al., 2008). These cells have distinctive functional characteristics that aid identification, including nonlinear spatial summation (Crook et al., 2008) and spatially segregated subunits within their receptive fields (Rhoades et al., 2019). In future work, we will record from this population and investigate whether these cells have different gain control signatures than the other M-layer neurons in our database, which are likely driven by parasol ganglion cells.

### Is separable gain control universal?

Our results in Figures **8** and **9** imply that the independence of luminance and contrast gain mechanisms reported by Mante et al. (2005) in the cat LGN also holds for M and P cells of the monkey LGN. Despite variability across and within cell types (see Fig. **4**), we could decompose temporal frequency tuning into changes driven solely by luminance or changes driven solely by contrast. The separability we observed supports the idea that gain control mechanisms reflect the independence of luminance and contrast observed in natural scene statistics (Frazor and Geisler, 2006).Evidence across species suggests that luminance adaptation of the kind we report here begins at the cones and propagates downstream (see review by Rieke and Rudd (2009)). Contrast adaptation involves changes at multiple sites of the inner plexiform layer of the retina (see Demb (2008) for a review) that are downstream of the cones. The current results show that little interaction between each adaptation mechanism is apparent within the primate LGN. The results imply that the LGN-specific adaptation mechanisms, initially reported in the primate by Kaplan et al. (1987), are uncorrelated with luminance adaptation. This separability of luminance and contrast adaptation is also a property of neurons in the cat’s visual cortex (Geisler et al., 2007). However, it is unknown whether this holds for primate V1. Future studies should explore sites of additional adaptation downstream of the primate LGN and establish whether this separability is a universal principle.

## Acknowledgements

We are grateful to Valerio Mante, Ethan Benardete, Bob Shapley, and Barry Lee for advice, feedback, and helpful discussions. This work was supported by NIH grants EY022428, EY007136, and EY013079 and by a grant from the Simons Foundation.

## Author contributions

RTR: design, methods, data collection, analysis, writing. JGK, JMH, PL: data collection. MJH: design, methods, writing. JAM: design, methods, writing.

